# Single molecule analysis reveals the role of regulatory light chains in fine-tuning skeletal myosin-II function

**DOI:** 10.1101/2020.01.21.913558

**Authors:** Arnab Nayak, Tianbang Wang, Peter Franz, Walter Steffen, Igor Chizhov, Georgios Tsiavaliaris, Mamta Amrute-Nayak

## Abstract

Myosin II is the main force generating motor during muscle contraction. Myosin II exists as different isoforms, involved in diverse physiological functions. The outstanding question is whether the myosin heavy chain (MHC) isoforms alone account for the distinct physiological properties. Unique sets of essential and regulatory light chains (RLCs) assembled with specific MHCs raises an interesting possibility of specialization of myosin functions via light chains (LCs). Here, we ask whether different RLCs contribute to the functional diversification. To investigate this, we generated chimeric motors by reconstituting MHC fast isoform (MyHC-IId) and slow isoform (MHC-I) with different light chain variants. As a result of RLCs swapping, actin filament sliding velocity increased by ∼ 10 fold for the slow myosin and decreased by >3 fold for the fast myosin. Ensemble molecule solution kinetics and single molecule optical trapping measurements provided in-depth insights into altered chemo mechanical properties of the myosin motors, thereby affecting the sliding speed. We find that both slow and fast myosins mechanical output is sensitive to the RLC isoform and propose that RLCs are crucial in fine-tuning of the myosin function.

## Introduction

Myosin motors drive diverse motile processes, ranging from intracellular cargo transport, cell division to muscle contraction and the whole cell movement. Striated muscle myosin such as skeletal or cardiac myosin 2, responsible for generating the force during muscle contraction is a hexameric motor composed of 2 heavy chains (MHC) and 4 light chains (LC) [1]. Each heavy chain contains a globular motor domain, alpha-helical light chain binding domain, and a long coiled-coil rod. The rod parts from different myosin molecules self-assemble to form bipolar thick filaments. The ∼9 nm long α-helix, also termed as ‘lever arm’ serves as a link between the motor domain to the rod part of heavy chains. Each MHC has an essential (ELC) and regulatory light chain (RLC) wrapped around the lever arm (cf. Figure 1A). The main motor domain has ATP and actin binding sites. During the chemo-mechanical coupling, small conformational change in the catalytic domain linked with the ATP hydrolysis gets amplified as a large movement of the lever arm [2, 3]. The light chains are reported to be essential to maintain the rigidity of the lever arm as force is transduced during the power stroke [4].

**Figure 1.**
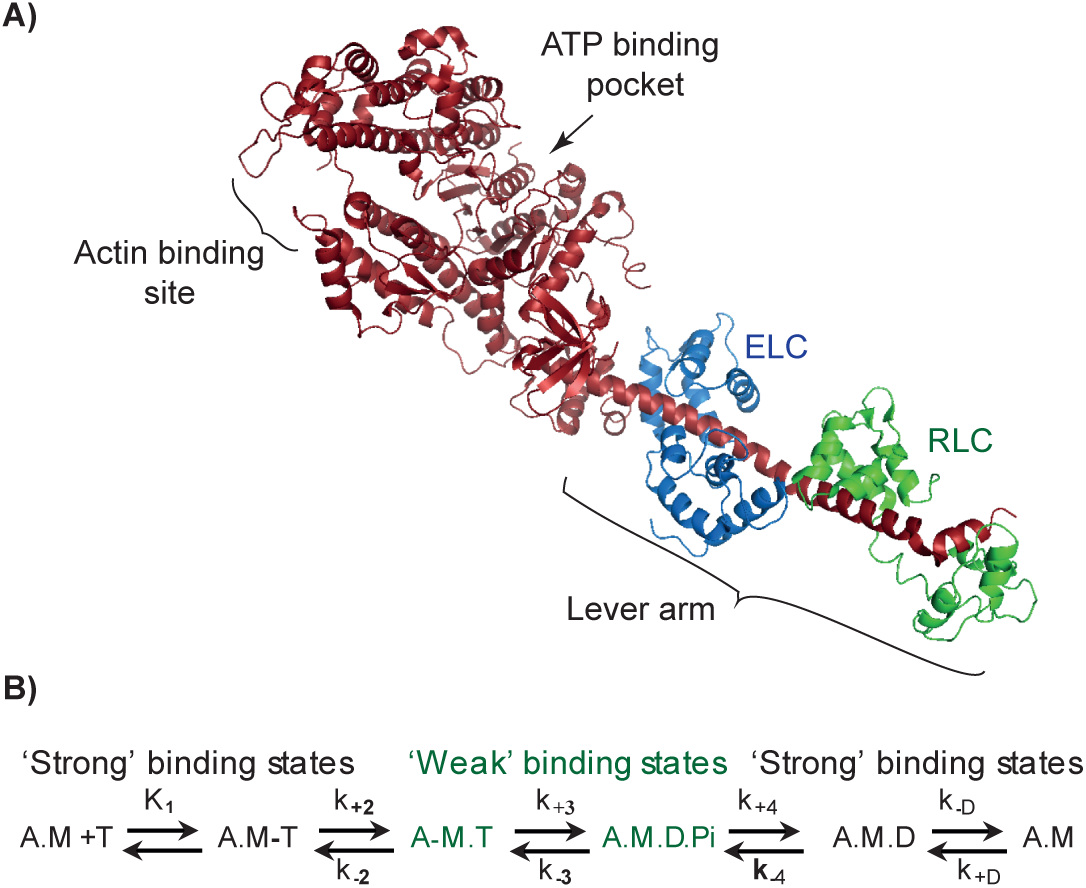
Myosin S1 structure and ATPase cycle. **A)** Crystal structure of scallop myosin II – subfragment -1, PDB-1SR6 [60]. The myosin heavy and light chains are shown as ribbon diagram; S1 is indicated with the key parts of the motor domain. **B)** Kinetic scheme for actomyosin ATPase. Myosin II with different nucleotide states are illustrated; A-actin, M - myosin, T - ATP, D - ADP, Pi - inorganic phosphate. Strong and weak interactions states of myosin with actin are labelled in black or green, respectively. Rate constants for forward and reverse reactions are referred to as, k_+n_ or k_-n_, respectively.

Muscle myosin II exists as different isoforms, three ‘fast’ myosin heavy chain (MHC) isoforms, i.e., MHC-IIa, -IId/IIx, and -IIb, and one ‘slow’ MHC-1 isoform [5]. MHC-1 isoform or beta cardiac isoform is primarily expressed in slow-skeletal, aerobic muscle fibers and in heart ventricle [6]. The muscle expressing ‘fast’-MHCs express myosin light chain (MLC) isoforms, i.e., different ratios of ELC isoforms, MLC1f and MLC3f, and RLC isoform, MLC2f. The ‘slow’-MHC-1 isoform is typically equipped with ELC isoform, MLC1s and RLC isoform, MLC2s/MLC2v.

RLC or MLC2 (∼19 kDa), encoded by the *MYL2*/*Myl2* genes, is a member of the EF-hand superfamily of Ca^2+^-binding proteins with a helix-loop-helix structural motif. MLC2 non-covalently binds to the distal end of the lever arm at the junction between S1 and S2 regions of myosin. While the C-terminal domain of RLC binds in the region between Glu 808 and Val 826 of myosin lever arm, the N-terminal domain of RLC surrounds the region between amino acids Asn 825 and Leu 842 [2]. Apart from its structural role to provide rigidity, RLC’s are known to regulate cardiac, smooth, and skeletal muscle contraction via phosphorylation. For example, genetic loss-of-function studies in mice revealed an essential function of RLCs in cardiac contraction [7]. Compelling reports demonstrated further that RLC phosphorylation regulates cardiac muscle contraction by increasing the number of cross-bridges available to generate the force [8–10]. Reduction in phosphorylation levels of MLC2v are critically linked with human cardiomyopathies [11]. Phosphorylation of MLC2 in smooth muscle myosin II determines the active state of the motor required for muscle contraction, thus making RLC a key regulator. However, for striated muscle myosin, the phosphorylation status of RLC is not correlated with the activation of the cross-bridge cycle, but rather Ca^2+^ binding to troponin activating the thin filaments is the prerequisite to initiate the muscle contraction. MLC2 phosphorylation is known to increase the steady-state force in permeabilized muscle fibers by increasing the number of cross-bridges interacting with actin filaments at submaximal levels of Ca^2+^ activation, without affecting maximum shortening velocity [12, 13]. Thus, RLCs exert differential effects in a tissue specific manner. Collectively, MLC-2 is thus considered to play a mechanical role by stabilizing the lever arm and a regulatory role through phosphorylation, but not to affect the ATPase kinetics.

Muscle fiber studies indicated that the cross-bridge kinetics underlying the force transients are determined mainly by the MHC isoforms [14]. While the MHCs as the main determinant of the mechanical output is well agreed on, MLC’s primary role in skeletal myosin II is considered to be rather structural. The correlation of the muscle fiber specific expression and assembly of the light chains with specific MHCs, however, might imply additional role for the regulatory light chains. Two major observations; 1) the distinct sets of light chains associate with different isoforms of myosin heavy chain, and 2) single point mutations in the ventricular RLC causing heart disorder in humans, such as hypertrophic and dilated cardiomyopathies, directed us to investigate the importance of RLC in mechanical performance of the myosins.

This investigation is also relevant to understand the mechanical performance of the hybrid/chimeric motors i.e., combination of a specific myosin heavy chains with different light chains, found under different physiological conditions. In some cases, heterogeneity in MHC and LC isoforms expression in single muscle fibers has been documented in vertebrate skeletal muscles. For instance, extensive analysis of the SDS-PAGE gels from single soleus muscle fibers revealed that in majority of fibers the slow heavy chain and corresponding light chains were expressed. In some soleus single fibers however, fast light chain isoforms were expressed, which potentially may form complex with the slow heavy chains [15]. In human fast single fibers co-existence of both fast (MLC2f) and slow (MLC2s) light chains was observed [16]. Moreover, analyses of single fiber segments revealed coexpression of fast- and slow-myosin components nonuniformly distributed along the fiber length. [17–19]. During developmental phase, the concentration of such hybrid/chimeric motors is reported to be increased [19]. Thus, a mixture of fast and slow motor components (i.e., heavy and light chains) indeed exists in single muscle fibers; however, the mechanical properties of such hybrid motors remain unexplored.

Here, we examined the myosin chimeras where the endogenous regulatory light chain was replaced with a different RLC variant to probe whether, how, and which biochemical and mechanical properties are affected as a result of different trimeric formation/s of myosin II motor complex (i.e., MHC, ELC, and RLC). We found that the motor properties of a given myosin isoform were sensitive to the type of RLCs. Chimeric motor based on fast myosin heavy chain translocated the actin filaments with reduced speed; conversely the slow myosin improved the gliding speed, simply by replacing native RLC with a different RLC variant. Steady state and transient ATPase kinetics measurements allowed us to relate the altered motor properties to changes in the overall ATP turnover time and rates of product release. Single-molecule analyses based on optical trapping measurements revealed pronounced alterations in the duration of the strong actin bound states, size of the power stroke, as well as the stiffness of motors. These studies unraveled the critical role of different RLC’s on muscle myosin motor function.

## Materials and methods

### Generation of single headed myosin motors from native myosin II

Full length rabbit fast skeletal muscle myosin II, MyHC-IId/, and MHC-I was isolated from skinned fibres of *M. psoas* and *M. soleus* muscle as previously described [20, 21]. The myosin II was digested with papain in order to generate single headed subfragment-1 (myosin S1), respectively, as described earlier [22, 23]. S1 was stored at -80°C in 50% glycerol in a buffer (5 mM Na-phosphate, 10 mM K-acetate, 4 mM Mg-acetate and, 2 mM DTT at pH 7.0). In the reconstitution experiments, freshly prepared S1 was employed for further downstream processing for light chain exchange.

To collect the muscle tissues, the rabbits were euthanized as per the guidelines from German animal protection act §4 (Tötung zu wissenschaftlichen Zwecken/ sacrifice for scientific purposes). The animal was registserd under reference number 2016/122, obtained from Medical School Hannover central animal facility department. All the procedures were carried out in accordance with relevant guidelines and regulations from Medical school Hannover, Germany. No experiments were performed on live animals prior to the sacrifice.

### Regulatory light chain expression

The plasmid vector (EX-T0572-B09) containing full length human cardiac, slow light chain 2 insert (i.e., *MYL2v*) and vector (EX-D0356-B09) containing human, fast skeletal muscle myosin regulatory light chain (*MYLPF*) sequence were purchased from Genecoepia (Rockville, MD, US).The plasmid vector containing BDTC (biotin-dependent transcarboxylase) chicken gizzard smooth muscle regulatory light chain (cgmRLC) was kindly provided by Dr. Atsuko Iwane to Walter Steffen. The details of BDTC-cgmRLC fusion protein are provided in Iwane et al., 1997[24]. As mentioned in the Iwane et. al., a recombinant fusion protein, BDTC-cgmRLC was generated by fusing biotin-dependent transcarboxylase (BDTC) with chicken gizzard muscle myosin RLC sequence at N-terminus. BDTC is a 123 amino acid peptide sequence of the 1.3S subunit of *Propionibacterium shermanii* transcarboxylase. BDTC was post-translationally biotinylated by biotin holoenzyme ligase in *Escherichia coli* (*E.coli*) *in vivo.* Biotin was added to the bacterial culture in growth medium after the induction of the bacterial cells with Isopropyl β-d-1-thiogalactopyranoside (IPTG). The molecular weight of the BDTC-cgmRLC protein is ∼34 kD.

The plasmids (EX-T0572-B09, EX-D0356-B09) were transformed in Rosetta^TM^ competent cells (EMD Millipore, Burlington, MA, USA) for expression of the light chains, MLC2v and MLC2B, respectively. The isolation and purification of the protein was followed as described previously [24, 23]. Please note that bacterially expressed regulatory light chains, BDTC-cgmRLC, MLC2B (or MLC2F), and MLC2v were without post-translational modifications such as phosphorylation.

### Regulatory light chain exchange

RLC exchange with S1 was performed as previously described [24,23,25–27], with some modifications. Myosins, S1f (psoas fast myosin subfragment 1) or S1s (soleus slow myosin subfragment 1) were incubated with ∼10 fold molar excess MLC2v/MLC2B/cgmRLC for 30 min at 30°C in a buffer (Mg^2+^ free condition) containing 50 mM HEPES (pH 7.6), 0.5 M NaCl, 10 mM EDTA, 10 mM DTT. The exchange reaction was stopped by the addition of 12 mM Mg^2+^ to the reaction mixture, and incubated on ice for 30 min. To remove the excess RLCs from the reconstituted myosin and to obtain the active motors, myosin was further incubated with high concentration of actin filaments for 30 min on ice. The mixture was centrifuged over 20 % sucrose cushion buffer at 70,000 rpm for 30 min. The pellet was once washed with buffer (5 mM Na-phosphate, 10 mM K-acetate, and 4 mM Mg-acetate, 2 mM DTT, pH 7.0) and then resuspended in the buffer with added 10 mM ATP to release active myosin from actin filaments. The actin filaments were separated from myosin solution by centrifugation. The supernatant containing myosin motors was mixed with 50% glycerol, flash frozen in liquid nitrogen and stored at -80°C, and used for further experiments as required. The probes were run on the 15 % polyacrylamide gels and stained with Coomassie stain (Phastgel Blue R-350 from GE healthcare, US). The gel images were acquired and either Image Quant or Image J program was used for the densitometric analysis to determine the RLC exchange efficiency for the chimeric myosins. To measure the intensities of the RLC bands, region of interest (ROI) (including both the native and exchanged RLC bands) was selected and plotted as intensity histogram. The area under the curve for each respective band was measured for the intensity values. The difference in the molecular weight of the protein was taken into account while calculating the exchange efficiency. About ≥85 % RLC exchange was observed for several preparations of S1 as shown in Figure S1. The myosin was used for motility and single molecule assays. For the solution kinetics experiments, the myosin chimeras were prepared without final actin co-sedimentation step to avoid ATP in the myosin solution. Instead, after the RLC exchange reaction was completed, the myosin was loaded on spin columns with high molecular weight cut-off (100 kDa) to separate the free RLCs from the assembled myosin complexes. The removal of free RLC was ensured by multiple dilutions with buffer and passing through the spin columns. The myosin in buffer (5 mM Na-phosphate, 10 mM K-acetate, and 4 mM Mg-acetate, 2 mM DTT, pH 7.0) containing 3 % sucrose was finally flash frozen and stored at -80°C.

### Preparation of actin filaments

Actin filaments were generated by incubating rabbit G-actin in polymerisation buffer containing 5 mM Na-phosphate, 10 mM K-acetate, and 4 mM Mg-acetate, supplemented with protease inhibitor overnight at 4°C. Equimolar concentration of fluorescent phalloidin was added to fluorescently mark the actin filaments. To get long biotinylated actin filaments (≥ 20 µm) for optical trapping experiments, additionally, 1 mM DTT and 1 mM ATP was added to the polymerisation mixture as described by Steffen et al. [28].

### Active myosin heads

Prior to use in *in vitro motility* assays or optical trapping measurements, the myosin was further purified to discard any inactive motors by co-sedimentation with concentrated F-actin. Actin-myosin complex was dissociated by addition of 4 mM Mg.ATP. While the active heads released from actin in the presence of ATP, the inactive myosin remained bound and separated by centrifugation at 70,000 rpm for 30 min at 4°C. The supernatant containing the enzymatically active motors was further supplemented with 2 mM DTT and protease inhibitor and used in the functional assays. This procedure to remove the inactive motor heads was routinely followed prior to the main experiments.

### Steady-State ATPase and transient kinetic experiments

Actin-activated ATPase was measured with a BioTek Synergy 4 multiplate reader (BioTek) using the nicotineamide adenine dinucleotide (NADH) coupled assay. A solution of myosin (0.2 µM), F-actin, NADH (0.4 mM), lactate dehydrogenase (LDH; 0.02 mg/ml), pyruvate kinase (PK; 0.05 mg/ml) and phosphoenolpyruvate (PEP; 0.5 mM) in Buffer containing 25 mM HEPES pH = 7.3, 25 mM KCl, 5 mM MgCl_2_, 0.5 mM DTT was mixed with ATP (2 mM) to start the reaction. ATPase rates at different concentrations of F-actin were obtained from the slopes of the corresponding linear fits of the time dependent absorbance change at 340 nm. Rates were plotted against the F-actin concentration and data were fitted according to Michaelis-Menten kinetics. Transient kinetic experiments were carried out with a Hi-Tech Scientific SF-61 DX double mixing stopped-flow system (0.5 ms dead time) at 20 °C. Experimental buffer contained 20 mM MOPS pH = 7.0, 25 mM KCl, 1mM DTT, 5 mM MgCl_2_. For ADP-release measurements a solution of 2 µM actomyosin (AM) preincubated with 20 µM mantADP was rapidly mixed with 1.6 mM ADP. The reaction was followed through a KV389 cut-off filter by monitoring mantADP fluorescence changes induced by fluorescence resonance energy transfer (FRET) through tryptophan excitation at 296 nm. P_i_-release was measured under single or multiple turnover conditions with ATP as substrate as described before [29].

### In vitro motility assay

*In vitro motility* assay was performed with monomeric S1 motors with native or exchanged RLC, by immobilisation of the motors on nitrocellulose coated surface. The assay is described in more details in [20]. Briefly, myosin was incubated for 5 min on nitrocellulose coated surface, followed by wash and surface blocking with 1 mg/ml BSA in assay buffer (AB; 25 mM imidazole hydrochloride pH-7.2, 25 mM NaCl, 4 mM MgCl2, 1 mM EGTA, and 2 mM DTT). To block the inactive or damaged myosin motor heads 0.25 μM short, unlabelled F-actin was injected in the flow cell and incubated for 1 min. 2 mM ATP was introduced in the chamber to release the actin filaments and to make the active motor heads accessible. ATP was washed out with AB buffer, TMR (tetra methyl rhodamine) labelled F-actin was incubated for 1 min, washed to remove excess filaments, and finally the chamber was infused with buffer containing 2 mM MgATP and antibleach system (18 µg/ml catalase, 0.1 mg/ml glucose oxidase, 10 mg/ml D-glucose, and 10 mM DTT) to initiate F-actin motility. Images were acquired with a time resolution of 200 ms (i.e., 5 frames/sec) using a custom-made objective-type TIRF microscope. Actin filament gliding speed was analysed with Manual Tracking plug-in from ImageJ.

### 3-bead assay with optical tweezers

For the assay, flow cells with approximately 15 µl chamber volumes were assembled using coverslip with nitrocellulose coated beads. Glass microspheres (1-1.5 µm) suspended in 0.1 % nitrocellulose in amyl acetate were applied to 18x18 mm coverslips. All the dilutions of biotin-actin filaments and myosin S1 were made in reaction buffer containing 25 mM KCl, 25 mM Hepes (pH 7.6), 4 mM MgCl_2_, and 1% β-mercaptoethanol. For the experiment, the chamber was prepared as follows, 1) flow cells were first incubated with 1 µg/ml myosin S1 for 1 min, 2) followed by wash with 1 mg/ml BSA and incubated further for 1 min to block the surface, 3) finally, reaction mixture containing 0.8-1µm neutravidin coated polystyrene beads and 1–2 nM biotinylated actin filaments [30] was flowed in with 10 µMATP, ATP regenerating system (0.01 mM creatine phosphate, and 0.01 unit creating kinase) and deoxygenating system (1 mg/ml catalase, 1 mg/ml glucose oxidase, 2 mg/ml glucose, and 1% β-mercaptoethanol). As shown in the schematic view in Figure 4A, pre-stretched, biotinylated actin filament was held between the two optically trapped beads via neutravidin-biotin link forming a dumbbell, such that the actin filament forms a connection between two beads. The approximate distance between the two trapped beads was 10 µm. Low-compliance links between the trapped beads and the filament were adjusted such that the ratio of the position variance during free and bound periods was 5 -10 as described in Smith et al.[31]. All the experiments were performed at room temperature of approximately 22°C. WT and chimeric motor S1 were immobilized directly on nitrocellulose coated surface to have comparable conditions for the single molecules. Actin dumbbell was brought in contact with the myosin bead, and the actomyosin (AM) interaction events were registered as a reduction in free Brownian noise of the 2 beads. The 2 bead positions were precisely detected with quadrant photodetector, recorded and analyzed.

Data traces were collected at a sampling rate of 10,000 Hz and filtered at 5000 Hz. To improve the time resolution and detect short-lived AM binding events, high-frequency triangular wave of ∼600 Hz was applied to one of the traps as described in Veigel et al. [32]. Matlab routines were employed to evaluate data records for ‘AM’ interaction lifetime, ‘*t*_on_’ and, stroke size of motors.

### Single myosin molecule interaction with actin filaments in optical trapping measurements

To ensure that each data record is derived from an intermittent interaction between single myosin head and actin, myosin density on the bead surface was adjusted by diluting myosin solution. In our measurements, typically, one bead was found to interact with dumbbell among 8-10 beads scanned for the presence of motor on the bead. This measure minimized the likelihood of multiple molecules simultaneously interacting with the actin filaments. From the binomial probability distribution, in our measurements chances of presence of 2 motor per bead are < 1 %, and ∼9 % with 1 molecule per bead. From a total of 126 beads we analyzed for AM binding events; statistically, 0.4 beads may have 2 motors. This estimation, however, does not take into account the motors that are inaccessible to actin filament due to the positional limitation on the bead. Therefore, the probability of multiple motors interacting simultaneously with an actin filament is further lower.

### Statistical analysis

Binomial probability distribution analysis was used to estimate the chances of more than 1 molecule interacting with actin filaments in optical trap measurements. Unpaired t-test was used to calculate the statistical differences in the gliding velocities, single motor stroke size, and stiffness of the native vs chimeric motors. The nonparametric Mann-Whitney U test was used to calculate the statistical differences in the duration of AM interaction events, *t*on for the WT and chimeric motors. The statistical tests used to calculate statistical significance are included in the manuscript at relevant sections.

All the experimental protocols were performed in accordance with guidelines for good scientific practices and approved by the Medical School Hannover, Germany.

## Results

### Reconstitution of chimeric myosin II motors, and analysis of actin filament gliding

We investigated the role of the RLCs in modulating the motor function of skeletal myosin II isoform found in the fast and slow twitch muscle fibers. We isolated myosin II motors from rabbit fast-twitch, *Musculus psoas,* and slow-twitch, *Musculus soleus* muscle fibers. We inspected the native fast and slow myosin extracted from respective muscles in the SDS-PAGE gels to ensure the purity (Figure S1). We generated single headed subfragment -1 (S1) and reconstituted the chimeric motors based on fast, MyHC-IId, and slow MHC-I myosin II. In its native form, MyHC-IId henceforth referred as wild type fast (WT-S1f) myosin S1 associates with the 2 light chains, i.e., essential light chain (ELC), MLC1f/MLC3f, and the RLC, MLC2B. MHC-I or wild type slow (WT-S1s) myosin S1 are typically equipped with MLC1v and MLC2v.

As shown in Figure 2A, we used 3 different RLCs, i.e., human fast skeletal regulatory light chain, MLC2B, human slow skeletal or ventricular RLC, MLC2v, and chicken gizzard smooth muscle myosin RLC (cgmRLC) to generate 6 different chimeric motors based on the fast and slow myosins. The cgmRLC was previously reconstituted with the chicken skeletal myosin II and was shown to retain the high gliding velocity of the motor [24]. For smooth muscle myosin (SMM) lower gliding speed from 0.2 to ∼1 µm/s were reported under different motility conditions [33–35]. SMM displayed similar kinetics properties to that of slow skeletal soleus myosin [6] with basal Mg^2+^ ATPase activity of 0.05 s^-1^, and Acto-S1 ATPase v_max_ of 0.7 s^-1^, ADP dissociation of 15 s^-1^(at 20°C), and a second order rate constant for ATP binding of 0.5 µM^-1^s^-1^ [36]. Despite these SMM properties, when it was assembled with chicken skeletal muscle myosin, the actin filament gliding speed of the myosin subfragment-1 was increased by ∼4 fold, similar to the native full length skeletal myosin [24].

**Figure 2.**
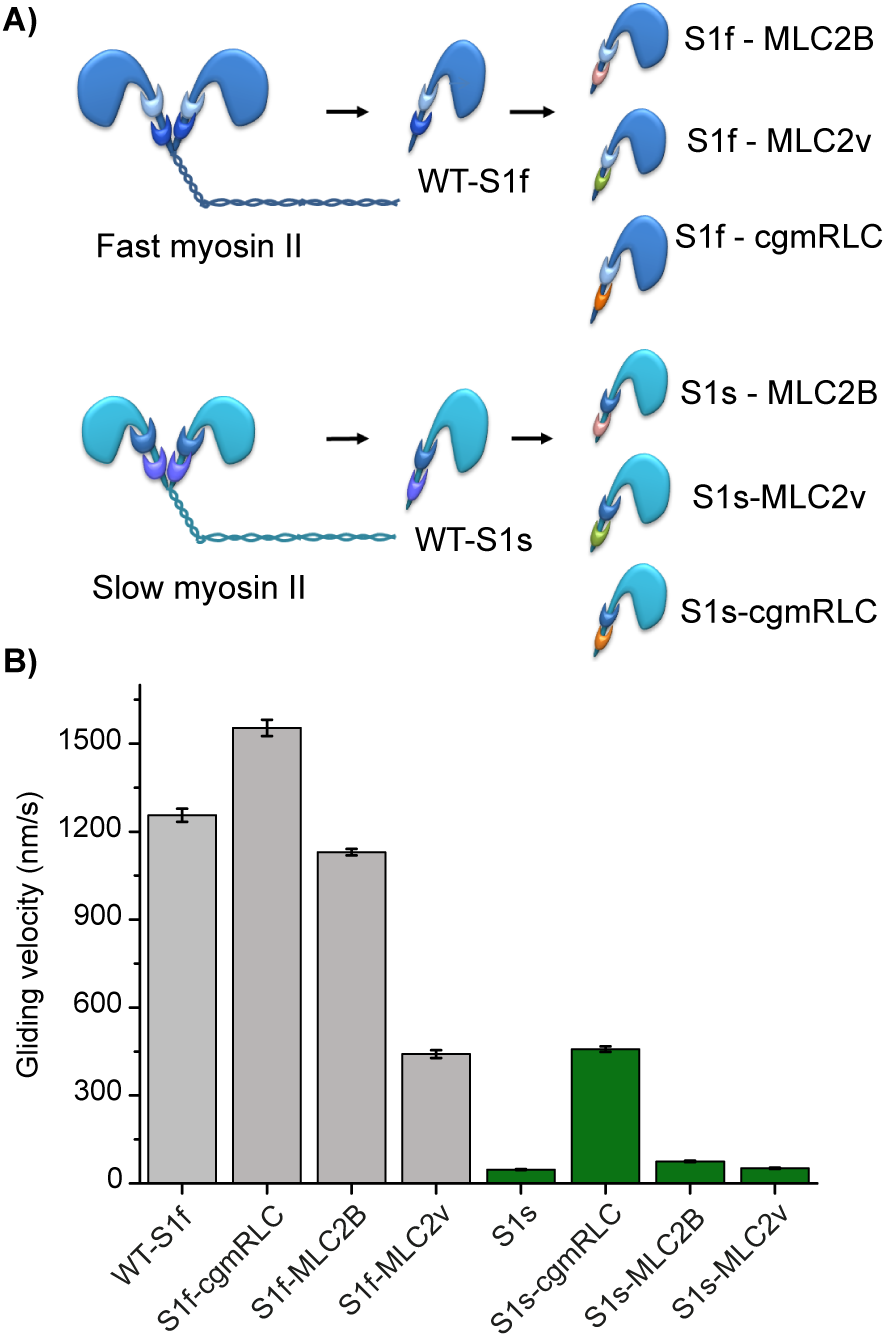
Reconstitution of chimeric motors. **A)** Scheme illustrates *in-vitro* reconstituted motors with different combinations of RLCs with fast (dark blue) and slow myosin II (sky blue) heavy chains. Papain digestion of full length myosin II generated single headed myosin S1. Regulatory light chains are color coded to indicate the exchange, MLC2B (pink), MLC2v (green) and cgmRLC (orange). **B)** *In vitro* motility assay using native myosin S1 (WT-S1f and WT-S1s) or with chimeric motors. Motors were immobilized on a nitrocellulose coated surface. Speed of movement was measured at saturating ATP concentration of 2 mM, at room temperature, 22°C. Bar diagrams show the reduction in mean velocity from 1.2 ± 0.19 for WT-S1f to 0.44 ± 0.13 µm/s for S1f-MLC2v (N = 72 and 100 actin filaments, respectively). Increase in mean velocity for WT-S1s from 0.047 ± 0.05 to 0.40 ± 0.05 µm/s for S1s-cgmRLC (N = 82 and 166 filaments, respectively). The motility experiments were performed with at least 3 different preparations of myosin motors and chimeras, and were highly reproducible. S1f-cgmRLC (N = 70), S1f-MLC2B (N = 110), S1s-MLC2v (N = 56), S1s-MLC2B (N = 55). Error bars represent ± SEM. Statistical significance calculated using unpaired t-test for the following pair of motors. WT-S1f and S1f-cgmRLC, P < 0.0001; WT-S1f and S1f-MLC2v, P < 0.0001; WT-S1s and S1s-cgmRLC, P < 0.0001; WT-S1s and S1s-MLC2B, P < 0.0001.

Therefore, we used it as a control to test its effect on the fast and slow skeletal myosin under our experimental conditions. As depicted in the Figure 2A, we replaced the endogenous RLCs of wild type myosins and generated following chimeric forms, 1) fast myosin chimeras - S1f-MLC2B (i.e., fast MHC with skeletal RLC), 2) S1f-MLC2v (i.e., fast MHC with ventricular RLC), 3) S1f-cgmRLC (i.e., fast MHC with smooth mucle RLC), and the slow myosin chimeras-4) S1s - MLC2v (i.e., slow MHC with ventricular RLC), 5) S1s-cgmRLC (i.e., slow MHC with smooth mucle RLC), and S1s-MLC2B (i.e., slow MHC with skeletal RLC). We employed human variants of the RLCs because of the nearly identical protein sequences as shown in Figure S2. Human MLC2B (hMLC2B) and rabbit MLC2f (rMLC2f) shares 98 % sequence similarity. Likewise, human MLC2v is 98% similar with the rabbit slow RLC isoform. However, the fast isoforms hMLC2B or rMLC2f has an 86 % sequence similarity with the slow MLC2v. The cgmRLC shares low sequence similarity with both fast MLC2B and slow MLC2v i.e., only 67 %.

To understand the main effect of RLC’s in these different combinations of chimeric motors, we focused on the mechanical parameters at the ensemble level using *in vitro* actin filament gliding assay

We characterized two-wild type and six-chimeric motors in *in vitro* actin filament gliding assays performed at 2 mM ATP concentrations (and ∼ 50 µg/ml surface density of motors). The motors were immobilized on nitrocellulose coated surface. As shown in Figure 2B and table 2, WT-S1f and WT-S1s showed >20 fold difference in the gliding speed i.e., 1.25 µm/s and 0.047 µm/s, respectively (cf. movie S1). Very interestingly, we found gliding velocities on both fast MyHC-IId and slow MyHC-I motor surface to be sensitive to the variant of RLCs bound to the lever arm. S1f-MLC2v combination resulted into decrease in the actin filament gliding velocity by ∼3 fold, while S1f-cgmRLC and S1f-hMLC2B retained the high actin filament gliding speed similar to the WT-S1f. On the contrary, S1s-cgmRLC chimeric motor showed ∼10 fold increase in the gliding speed of the slow S1s motor from 0.047 µm/s to 0.43 µm/s. S1s-MLC2B motor showed a marginal increase in the gliding speed compared to WT S1s. Interestingly, as shown in movie S2 and Figure 2B, fast and slow myosin driven actin filaments were indistinguishable when fast MHC was combined with MLC2v (S1f-MLC2v) and slow MHC with cgmRLC (S1s-cgmRLC).

These experiments showed that the motile activity of skeletal muscle myosin II is critically influenced by the regulatory light chain isoform. Replacement of the native RLCs from fast and slow myosin II S1 with the RLC variants, MLC2B and MLC2v respectively, showed comparable sliding velocities (Figure 2B and table 2). These results ruled out the alteration of motor properties due to the exchange procedure, and further emphasized RLC specific effects on myosin function.

The altered sliding velocities indicated that RLC interferes with the AM chemo-mechanical coupling. Actin filament sliding velocities are governed by the net rate of actomyosin cross-bridge cycling [37, 38]. The parameters that influence the sliding speed (*V*) include: (i) the lifetime of the strong AM bound state (*t*_on_), and (ii) the stroke size (*d*) of myosin, i.e., *V* = *d*/*t*_on_. Duration of AM bound and unbound states during the ATPase cycle described by duty ratio of the myosin, is the fraction of the total ATPase cycle time myosin spends bound to actin. The duty ratio can change either by a change in the ADP release rate from the AM complex or the rate of weak-to-strong bound transition of AM (i.e., A.M. D.Pi to A.M.D state); cf. ATPase scheme in Figure 1. One other parameter that must be considered in ensemble molecule sliding velocity measurements is the surface density of the myosin molecules. The skeletal myosin II, a low duty ratio motor (< 0.05), however showed low dependence of velocities on the motor density [39, 40].Therefore, we presumed that the velocities most likely changed due to altered *t*_on_ and/or *d*.

For native S1f, while some reports showed an isomerization step before Pi release as a rate limiting step, the others suggested Pi release as the slowest step in the AM ATPase cycle [41, 42] and ADP release to be fast (>500 s^-1^). For the WT S1s however, ADP was shown to have ˃10 fold higher affinity, and thus slower ADP release during the cross-bridge cycle, ∼ 20 s^-1^[6]. Therefore, the detachment rate of the AM complex at the postpower stroke state is limited by either rate of ADP release or ATP binding to the rigor AM complex for these slow and fast myosin isoforms.

### Ensemble molecule steady state ATPase and transient kinetics

To investigate the cause of the altered sliding velocities, we measured actin-activated ATPase activities and rates of product release to determine potential rate limiting steps in the ATPase cycle (Fig. 3). Consistent with previous studies [6] we obtained comparable basal ATPase activities between WT-S1f (0.04 s^-1^) and S1s (0.05 s^-1^) and almost 100-fold (3.5 s^-1^) and 10-fold (0.5 s^-1^) increase in the actin-activated ATPase activities, respectively (Fig. 3A, B, and Table 1). S1f-cgmRLC displayed comparable actin-activated ATPase activity as the wild-type construct (Table 1), whereas S1fMLC2v showed reduced actin activation as previously reported [27] (Table 1). For the soleus constructs we observed pronounced differences in the basal ATPase activity by almost a factor of 10 between S1s (0.05 s^-1^) and S1s-cgmRLC (0.55 s^-1^). The high basal ATPase activity of S1s-cgmRLC was unexpected but reproducibly observed with different preparations. Such comparably high basal activities have been observed in mutant myosins [43, 44], and also unconventional myosins Myosin IX [45], Myosin VIIa [46] displayed similar high basal activities. The S1sMLC2B construct showed a similar basal activity as the wild-type S1s (0.04 s^-1^). The ATPase activity increased for all constructs with increasing actin concentrations yielding values for activation at 60 µM F-actin of 0.14 s^-1^, 0.62 s^-1^ and 0.09 s^-1^ for S1s, S1s-cgmRLC, and S1s-MLC2B, respectively. Notably, the efficiency by which actin activated the ATPase (k_cat_/K_app_) was changed from three to five-fold among the motors. (Table 1). In additional transient kinetic experiments we measured the rates of ADP and Pi release as major determining factors of ATPase and sliding velocity. Generally, ADP release from actin-bound S1f is a very fast process with an estimated rate constant >500 s^-1^ [6]. As seen in Fig. 3C, ADP-release kinetics was too fast to be resolved or not accompanied by signal change. The amplitude change was limited by the dead-time (0.5 ms) of the instrument, providing only estimates for the rates of ADP release from acto-myosin of >500 s^-1^ for all constructs (Table 1). Notably, ADP-release from actin-bound S1s, could be monitored due to its slower release kinetics and high ADP-affinity [6]. We observed two-step ADP-release kinetics for S1s, S1s-cgmRLC, and S1s-MLC2B, yielding rate constants of k’_-AD_ = 54 s^-1^, 61 s^-1^ and 60 s^-1^ for the fast phase and k’’_-AD_ = 3.9 s^-1^, 4.9 s^-1^, and 2.9 s^-1^ for slow phase, respectively (Fig. 3D). We observed Pi-release as the rate-limiting step of the ATPase reaction for all measured constructs (Figure 3E and F). This is reflected in the similar rates between Pi-release rates and actin-activated ATPase rate at the 20 µM actin concentration (Fig 3A, B, Table 1). Thus, the RLCs appear to dictate the overall cycling time, mainly by affecting ADP release and modulating the Pi-release kinetics.

**Table 1.** Summary of kinetic parameters. *k*_basal_: ATPase activity in the absence of actin; *v_max_* maximum actin-activated ATPase activity; *K_app_*: concentration of actin at ½ *v_max_*; *v_max_ and K_app_* - estimated from hyperbolic fits; *v_max_* (60µM) - actin activated ATPase rate at 60 µM F-actin; *k*_cat_/*K*_app_: catalytic efficiency; *k*^fast^_-AD_, *k*^slow^_-AD_: rate constants of the ADP release from acto-myosin; *k*_-Pi_: rate constant of P_i_-release at 20 µM F-actin concentration.

|  |  | **S1f** | **S1f**<br>**cgmRLC** | **S1f**<br>**MLC2v** | **S1s** | **S1s**<br>**cgmRLC** | **S1s**<br>**MLC2B** |
| --- | --- | --- | --- | --- | --- | --- | --- |
| $k_{\text{basal}}$ | (s<sup>-1</sup>) | **0.04 ± 0.01** | **0.06 ± 0.01** | **0.04 ± 0.01** | **0.05 ± 0.01** | **0.55 ± 0.01** | **0.04 ± 0.01** |
| $v_{\text{max}} \text{ (fit)}$ | (s<sup>-1</sup>) | **3.5 ± 1** | **2.5 ± 0.1** | **1.0 ± 0.3** | **0.5 ± 0.2** | **0.6 ± 0.1** | **0.1 ± 0.1** |
| $v_{\text{max}}$<br>(60μM) | (s<sup>-1</sup>) | **0.66 ± 0.01** | **0.52 ± 0.2** | **0.30 ± 0.02** | **0.14 ± 0.01** | **0.62 ± 0.01** | **0.09 ± 0.01** |
| $K_{\text{app}} \text{ (fit)}$ | (μM) | **278 ± 96** | **244 ± 10** | **142 ± 54** | **208 ± 91** | **22 ± 3** | **39 ± 4** |
| $k_{\text{cat}}/K_{\text{app}}$ | (10<sup>-3</sup>μM<sup>-1</sup>s<sup>-1</sup>) | **10** | **11** | **4** | **2** | **4** | **2** |
| $k_{\text{-AD}}^{\text{fast}}$ | (s<sup>-1</sup>) | **> 500** | **> 500** | **> 500** | **54 ± 3** | **61 ± 1** | **60 ± 3** |
| $k_{\text{-AD}}^{\text{slow}}$ | (s<sup>-1</sup>) | **-** | **-** | **-** | **4 ± 1** | **5 ± 1** | **3 ± 1** |
| $k_{\text{-Pi}}$<br>(20μM) | (s<sup>-1</sup>) | **0.18 ± 0.01** | **0.29 ± 0.01** | **0.07 ± 0.01** | **0.05 ± 0.01** | **0.17 ± 0.01** | **0.03 ± 0.01** |

**Table 2.**
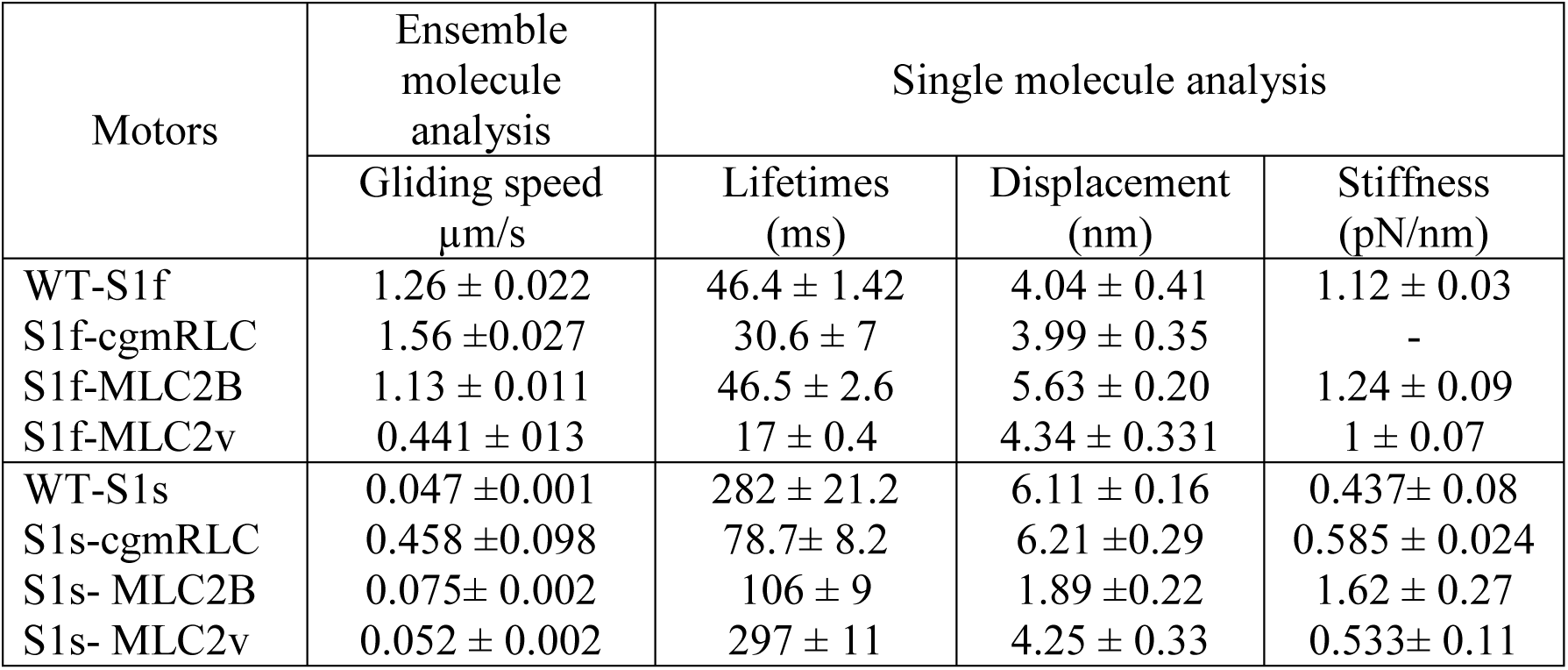
The measured parameters from ensemble molecule motility assay and single molecule optical trapping measurements are summarized in the table: actin filament gliding speed, lifetimes of AM bound state, displacement of actin by individual motors, and stiffness for wild type and chimeric motor forms.

**Figure 3:**
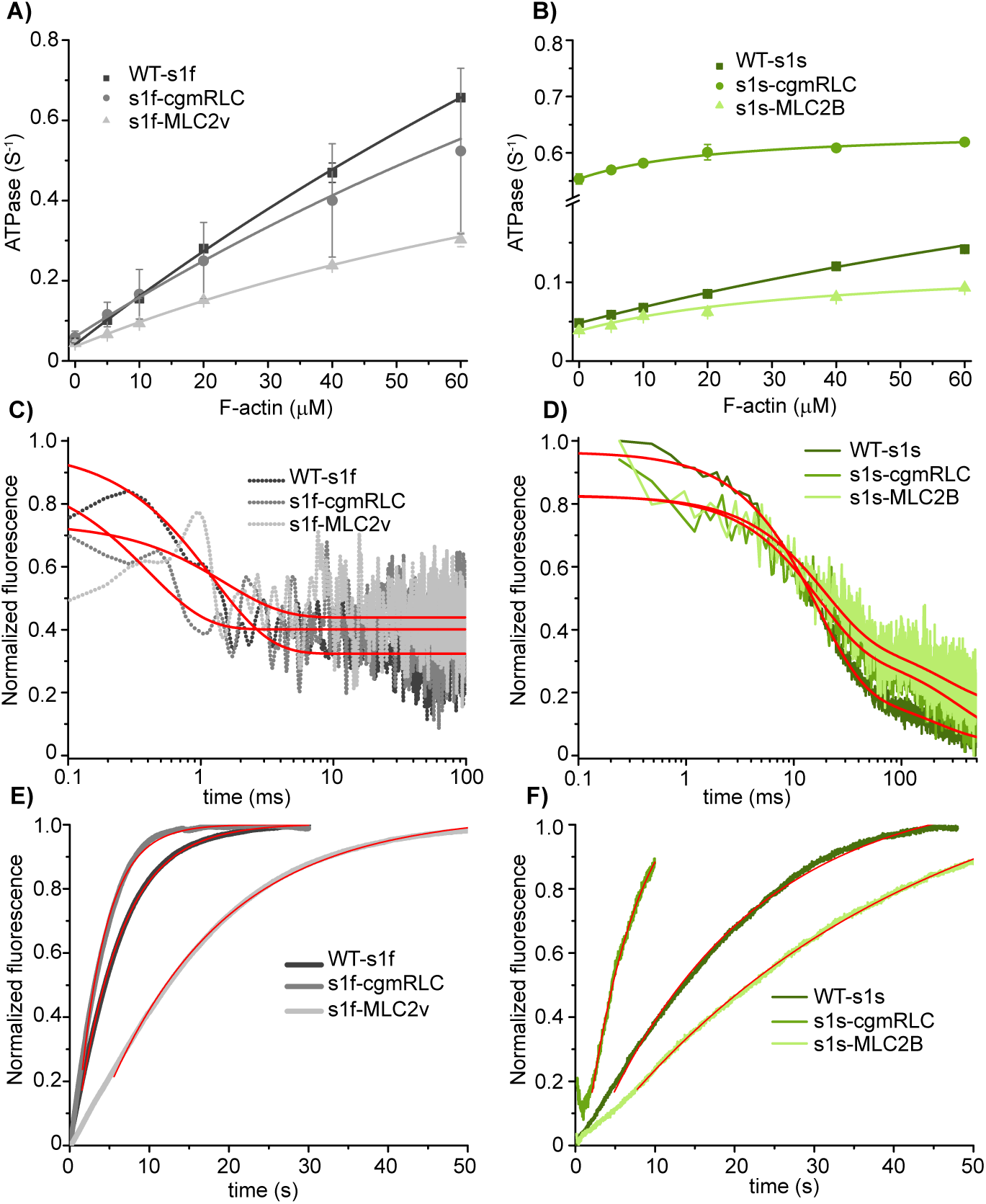
Ensemble kinetic experiments. Plots of ATP turnover rates as a function of F-actin concentration for fast (**A**) and slow (**B**) myosins. Data fitted to hyperbolas using the Michaelis-Menten formula. The catalytic efficiency kcat/Kapp, was obtained from the initial slope of the hyperbola. (C and D) ADP-release kinetics from acto-myosin. The rate of ADP dissociation from the acto-myosin complex was measured by rapidly mixing acto-myosin-mantADP complex with excess ADP. The obtained decrease in mant fluorescence followed in the case of the fast myosins single exponential kinetics (**C**). For the slow myosin constructs the fluorescence transients were best described by two exponentials with a fast rate constant (*k*^fast^-AD) and a slow rate constant (*k*^slow^-AD) indicative of two-step ADP-release kinetics (**D**). ATP turnover experiments of acto-myosin constructs using coumarin-labelled phosphate binding protein (MDCC-PBP, ThermoFisherScientific) as sensor for the detection of liberated Pi from fast (**E**) and slow (**F**) myosins. The transients follow single exponentials with corresponding rate constant for Pi release (*k*_-Pi_). All kinetic parameters are summarized in Table 1.

**Figure 4:**
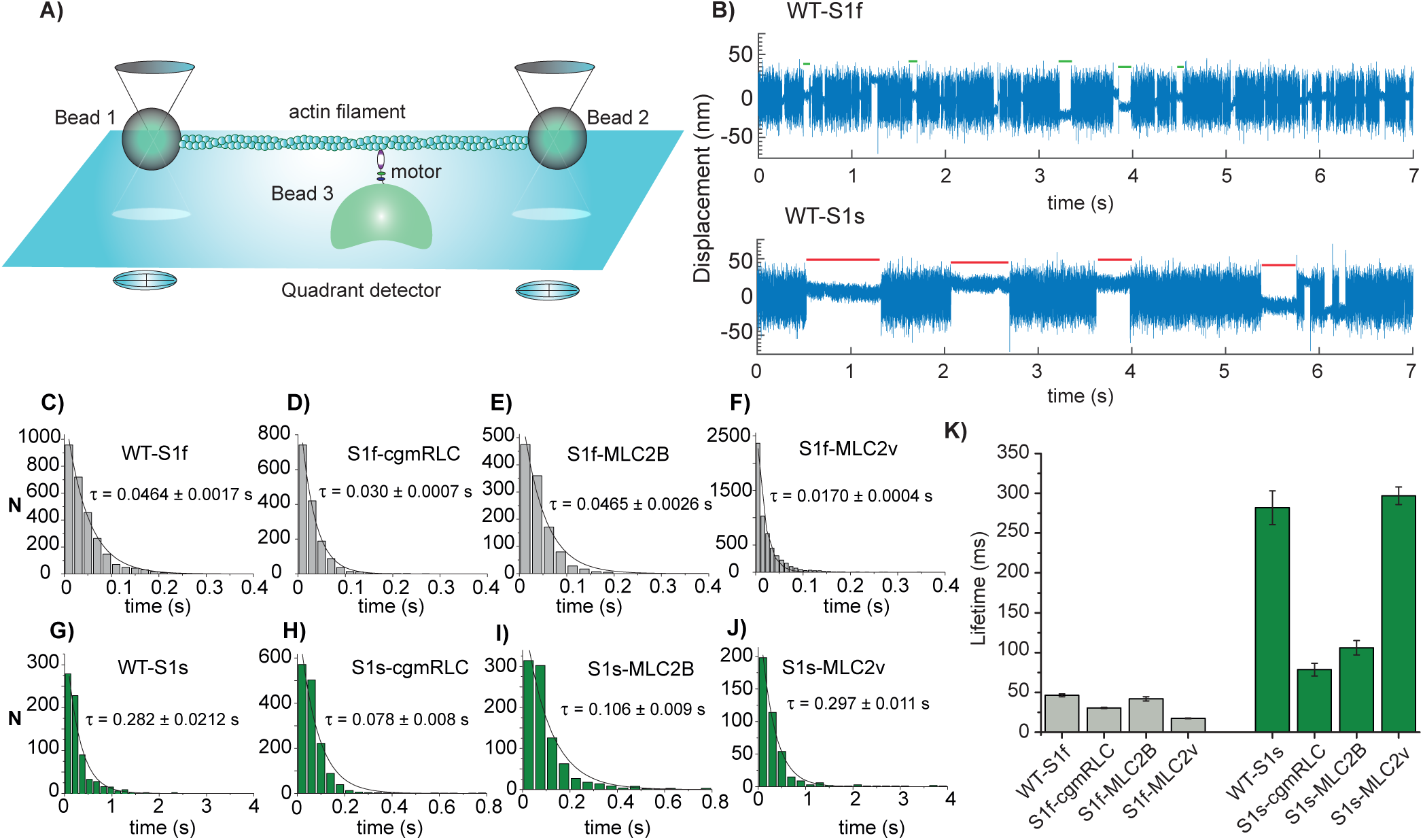
Single molecule optical trapping - lifetime of AM interactions. **A)** The experimental set up for 3-bead optical trapping measurements. Please note that the indicated components, bead size, and protein dimensions are not to the scale. **B)** Original displacement over time data records; actin - myosin interaction can be observed as reduction in the large thermal fluctuations from a bead at 10 µM ATP. The upper and lower panel shows records collected from fast and slow WT myosin S1, respectively. Some of the interaction events are shown as green (fast WT) and red (slow WT) lines to indicate the duration of lifetimes, t_on_. Measurements were done by applying positive feedback and triangular wave (∼600Hz). **C-J)** S1f and S1s motors and chimeras - t_on_ from below indicated number of events plotted in histograms and fitted with single exponential decay function to determine the average time constant **K)** The bar diagram shows the average *t*on for wild type and chimeric motors at 10 µM ATP concentration, and room temperature of ∼22°C. Error bars represent ± SEM (from the fits). For WT-S1f; n = 5000 events, N = 21, for WT-S1s; n = 1675 events, N = 19, for S1f-cgmRLC; n = 1517 events, N = 10, for S1f-hMLC2B; n = 1953 events, N = 16, for S1f-MLC2v; n = 5714 events, N = 22, for S1s-cgmRLC; n = 1468 events, N = 12, for S1s-MLC2v; n = 400 events, N = 4, for S1s-MLC2B; n = 900 events, N = 8. N = number of individual motor molecules, n = number of AM interaction events. Statistical significance was calculated for following pairs of motors using nonparametric Mann-Whitney U test. WT-S1f and WT-S1s, P < 0.00001; WT-S1s and S1s-cgmRLC, P < 0.0001; WT-S1s and S1s-MLC2B, P < 0.0001; WT-S1f and S1f-hMLC2B, P = 0.0691; WT-S1f and S1f-MLC2v, P < 0.0001, WT-S1f and S1f-cgmRLC, P = 0.114; WT-S1s and S1s-MLC2v, P = 0.0941. P <0.05 – statistically significant, P >0.05 –not statistically significantly different.

### Single molecule analysis

To further probe the change in velocities with chimeric motors, we employed single molecule analysis method to gain precise insights into the biochemical properties such as duration of AM strong bound states, and biophysical properties such as stroke size (*d*) and stiffness of native and chimeric motor forms.

### Single chimeric motor molecule’s kinetic properties

We employed 3-bead trapping assays for in-depth analysis of the biochemical and mechanical properties that led to either improved or decrease in the actin filament gliding speed driven by chimeric motors. Optical trap experimental set up has been described previously in detail [28]. Briefly, the actin filament is embedded and stretched between the two optically trapped beads forming a dumbbell, and the motor is immobilized on a glass bead attached on the chamber surface (Figure 4A). The positions of the 2-trapped beads are precisely monitored using quadrant photodiode and acquired.

Single headed wild type or chimeric motor molecules were analyzed for their intermittent interaction with the actin filaments as shown in Figure 4B at 10 µM ATP concentration. We determined the lifetimes of the motor interaction with actin. The reduced Brownian noise in the data traces were characteristic of the AM bound states, ‘*t*_on_’, whereas free dumbbell noise, ‘t_off_’ represented myosin detached states. It is well known that actin accelerates the Pi release, and Pi release from myosin active site is closely associated with the powerstroke generation. However, there is no clear consensus on whether Pi is released before, during or after the powerstroke. In our experimental set up, the initiation of acto-myosin association seen as the reduction in the noise amplitude of the bead coincides with the powerstroke generation.

Thus, ‘*t*_on_’comprised the post-powerstroke states with ADP present in the active site (AM.ADP) plus the time the active site is free of nucleotide (rigor state) until a new ATP-molecule binds and initiates the rapid detachment of the myosin head from the actin filament. This argument is valid if the Pi is released before or during the powerstroke. Some recent studies have supported the model that the powerstroke precedes the Pi release from the active site [47, 48]. However, to consider AM.ADP.Pi state as ‘strong bound state’ - contributing to the lifetime of attached event - duration of this state should be sufficiently long after the powerstroke (at least 2 ms i.e., the detection limit with our current experimental set up). We cannot distinguish between strong bound AM.ADP.Pi and AM.ADP states.

The AM detached state, *t*_off_, included the ATP hydrolysis and M.ADP.Pi state prior to association with actin (cf. ATPase scheme in Figure 1). As the ADP release from the active site is in the range of ˃500 s^-1^, we believe that for the fast myosins, primarily AM bound rigor state is acquired. At saturating ATP-concentrations the duration of the rigor state becomes too short, which is essentially undetectible for fast myosins. Therefore, for comparison among different native and chimeric motors we measured lifetimes, ‘*t*_on_’ at 10 µM ATP concentrations. Figure 4B shows the comparison of 7 s traces obtained with WT-S1f and WT-S1s, containing several distinct single AM binding events at 10 µM ATP. The obvious difference in the two traces with the two types of motors was the short *vs.* long lifetime of AM bound events. Consistent with previous observations, S1f showed shorter *t*_on_ as compared to S1s [49]. We estimated the average lifetime of AM bound states by fitting the ‘*t*_on_’ events from several molecules with a single exponential decay function (Figure 4C-J). Figure 4K and table 2 displays the average lifetimes for wild type and the chimeric motors. Duration of AM bound states for WT-S1f, S1f-cgmRLC, and S1f-MLC2B showed no significant difference. However, the S1f-MLC2v motor showed ˃2 fold decrease in *t*_on_. For slow myosin, compared to WT-S1s, S1s-MLC2v retained the extended *t*_on_. Interestingly, with S1s-cgmRLC the *t*_on_ was significantly reduced by ∼4 fold, and by ∼3 fold for S1s-MLC2B. From the reciprocal of average time constants we estimated the AM detachment rate constants and thereby the second-order rate constants of ATP binding (*k*_ATP_) for fast myosins, *k*ATP = 2.2 µM^-1^s^-1^ for WT-S1f, 3.26 µM^-1^s^-1^ for S1f-cgmRLC, 2.2 µM^-1^s^-1^ for S1f-MLC2B, 5.9 µM^-1^s^-1^ for S1f-MLC2v. Our estimation of *k*ATP for WT-S1f is consistent with previous studies [50, 51].

For all the chimeric constructs except for one (i.e., S1f-MLC2v), the increase in the duration of AM bound state associated with the decrease in the actin filament sliding velocity. The chimeric motors, S1f-cgmRLC and S1f-MLC2B showing comparable *t*_on_ (or *k*_ATP_) to WT-S1f, also displayed similar gliding velocities (cf. figure 2B). Similar tendency was observed WT-S1s and S1s-MLC2v.

### Mechanical properties

#### Stroke Size

Previously, single molecule analysis of motors showed that the lever arm length affects the amplitude of displacement (i.e., stroke size, *δ*) [52]. As an accessory protein associated with the lever arm, whether light chain influences stroke size is not explored.

We therefore investigated whether different variants of the light chains influenced the amplitude of stroke size of the chimeric motors. We determined the average stroke size from the shift of histogram method, introduced by Molloy et. al. [53]. As shown in Figure 5A-I and table 2, we noted significant difference in the mean displacement for fast and slow motors i.e., from 4.04 ± 0.41 nm to 6.11 ± 0.16 nm, respectively. Thus, consistent with previous reports for rat fast and slow skeletal myosin II-S1 [49], the average working stroke for WT-S1f was smaller than for WT-S1s. We found moderate yet significant increase in the stroke size for chimeric construct S1f-hMLC2B as compared to its wild type counterpart. For slow myosin heavy chain however S1s-MLC2v and S1s-MLC2B combination resulted into significant decrease of the average displacement amplitude, with S1s-MLC2B displaying merely 1.86 nm stroke size.

**Figure 5:**
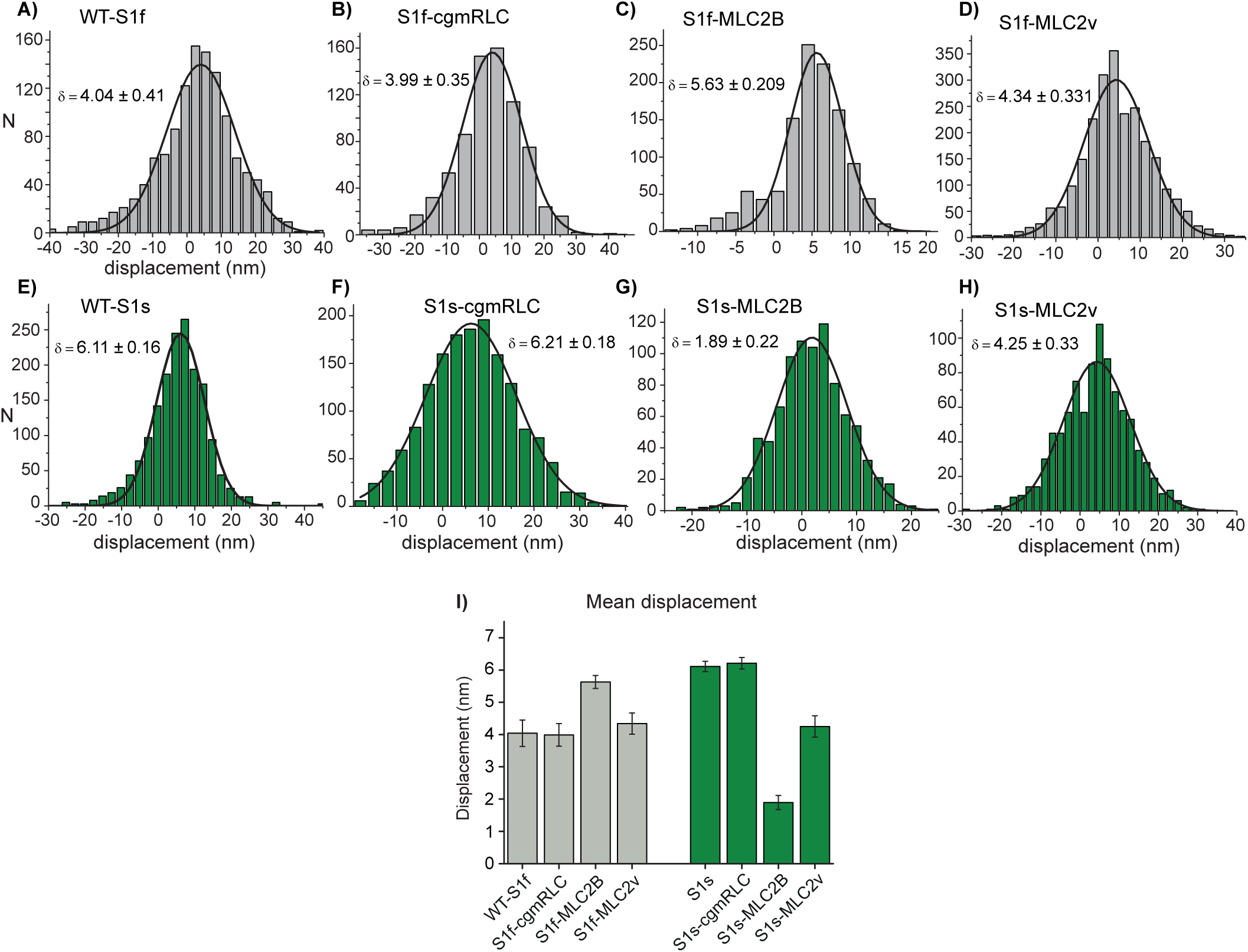
Powerstroke size of chimeric motors. **A-H)** The average stroke size/ mean displacement was determined by histogram shift (δ) from mean free dumbbell noise. The histograms fitted with Gaussian function to determine the average stroke size for indicated myosin motors. **I)** The average stroke size determined from the Gaussian fits for wild type and chimeric motors were compared in bar diagram. Error bars - standard error of mean (SEM from the fits). The statistical difference in the powerstroke size was calculated for the following pairs of motors. WT-S1f and WT-S1s, P < 0.0001; WT-S1f and S1f-cgmRLC, P = 0.8841; S1f and S1f-hMLC2B, P = 0.0053; S1f and S1f-MLC2v, P = 0.0025; WT-S1s and S1s-cgmRLC, P = 0.1002; WT-S1s and S1s-MLC2B, P < 0.0001; WT-S1s and S1s-MLC2v, P < 0.0001. Statistical significance calculated using unpaired t-test.

#### Stiffness

The light chains are reported to be responsible for the structural stability of the lever arm. The ∼9 nm long helical structure of lever arm that links the main motor core to the dimerization region and continued as a coiled coil region that forms the basis of the thick filament backbone may contribute to the compliance of the motor.

One assumption is that variants of RLCs provide different stiffness to the myosin head. Consequently, different myosin head stiffness may result in a different mechanical strain. The kinetics of some steps of the ATPase cross-bridge cycle is assumed to respond to the change of strain under load [54, 55].The strain dependence of ATPase kinetics is well reported for the motors such as myosin V and smooth muscle myosin [56, 36]. Could the binding of MLC2v in comparison to MLC2B change for example the stiffness of myosin S1f and resultantly alter the AM cross-bridge cycle?

To investigate this notion whether the different RLCs influenced the stiffness of the chimeric motors, we analyzed the AM interaction records acquired by applying positive feedback on the laser-trapped beads. Using bead variance-covariance method as described in [57], the stiffness of individual motors was calculated as shown in Figure 6. We found significant difference in the stiffness for WT-S1f *vs.* WT-S1s, i.e., 1.12 ± 0.03pN/nm and 0.437± 0.08 pN/nm, respectively (Figure 6E). The stiffness values are comparable with previous optical trapping measurements with rat fast and slow myosin S1[49]. For the fast myosin, replacing the light chains with 2 different variants (MLC2B and MLC2v) did not alter the stiffness significantly as shown in Figure 6 and Table 2. Similarly, for slow myosin chimeras (S1s-cgmRLC, S1s-MLC2v), there was slight increase in the average stiffness, however not significantly different from the wild type. S1s-MLC2B motor, however showed highest stiffness of 1.62± 0.27pN/nm.

**Figure 6.**
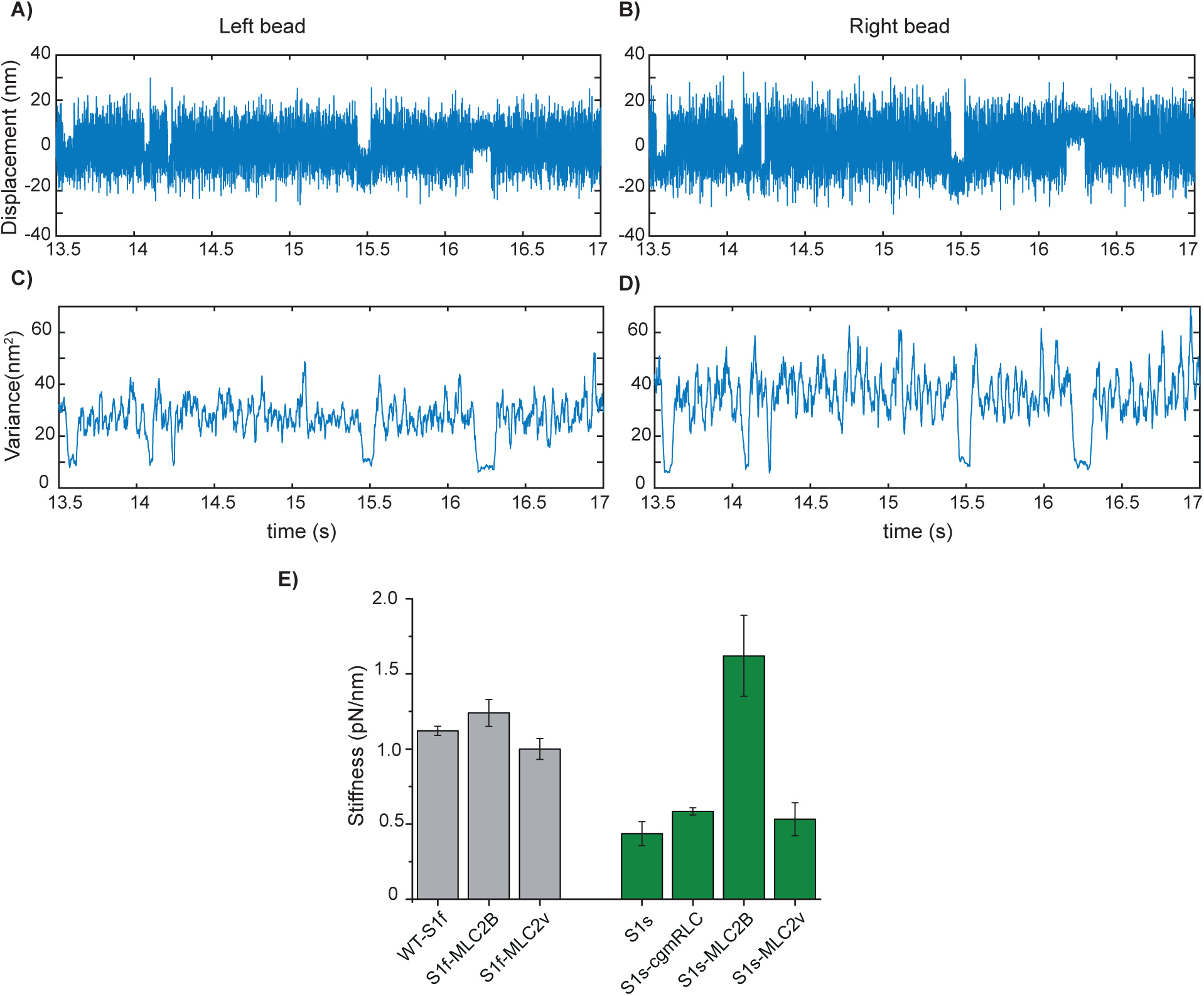
**Stiffness- A)** and **B)** original data records acquired in optical trapping experiments. Bead position is plotted over time for both left and right bead of the dumbbell. Positive position-feedback was used to increase the amplitude of thermal fluctuations, which effectively increase the variance ratio between binding events and free dumbbell noise for both traps in the direction of the actin filament axis. **C)** and **D)** variance versus time of the records shown in A and B. Variance was calculated for rolling window of 20 ms, at our sampling rate of 10 kHz. For the example shown here the variance-Hidden-Markov-method yielded a combined trap stiffness of 0.078 pN/nm and a myosin head stiffness of 0.85 pN/nm. **E)** Bar diagram with the stiffness measured for WT-S1f, WT-S1s and chimeric motors. Error bars - SD. Unpaired t-test was used to calculate the statistical significance. WT-S1f (N = 20) and WT-S1s (N = 13) displayed statistically significant difference with P < 0.0001. No statistically significant difference between WT-S1f and S1f-hMLC2B (N = 12), P = 0.72 or WT-S1f and S1f-MLC2v (N = 15), P = 0.128 was found. WT-S1s and S1s-cgmRLC (N = 10), marginal but statistically significant, P = 0.0493, WT-S1s and S1s-hMLC2B (N=6) P < 0.0001. For WT-S1s and S1s-MLC2v (N = 11), P = 0.4735. N = number of individual motor molecules.

To measure the stiffness of the WT-S1s and S1s-MLC2v chimera, the experiments were performed at higher ATP concentration i.e., at 50 µM ATP to avoid the occasional inclusion of contaminant fast myosin in the measurements. Although the soleus muscle primarily contains (about 95%) slow myosin II isoform, the presence of ∼5 % fast myosin isoform cannot be avoided. In ensemble molecule gliding assays, 5% contaminant motors is unlikely to influence the actin filament speed as previously measured for mixture of different ratios of myosin isoforms [35, 34]. Accordingly, more than 20 % contaminant motors were required to cause alteration of gliding speed. In the single molecule assays however, this factor needs to be cautiously controlled. At 50 µM ATP concentration the AM interaction events for the fast myosin are too short for detection with our set-up (<2 ms AM lifetime events could not be observed). This way we excluded the measurements from contaminant fast myosins. For S1s-cgmRLC, and S1sMLC2B stiffness was measured at 10 µM ATP concentration, as these motors showed ∼3 fold shorter t_on_ as compared to the WT-S1s and S1s-MLC2v, and ∼2 fold longer average t_on_ in comparison to the S1f chimeras.

Altogether, the mechanics measurement suggests that the light chains caused substantial alteration of the amplitude of powerstroke size and the motor stiffness for S1s-MLC2B, while for the other chimeras the parameters were either moderately altered or comparable with the wild type forms (Table 2).

## Discussion

For over 2 decades the issue of myosin heterogeneity in the muscle fibre has been discussed in the literature and particular emphasis has been put on the mixed content of myosin isoforms. However, until now it was difficult to systematically examine the mechanical performance of myosins associating with different types of RLCs. Here, by reconstituting homogenous population of such well-defined hybrid/chimeric motors and analysing the kinetic and mechanical features of the constructs using single molecule and ensemble measurements, we were able to unravel the influence of individual RLCs on the mechanical performance of slow and fast muscle myosins.

By swapping the RLCs between fast and slow myosin heavy chain isoforms, we showed that MLC2 naturally associating with fast myosin motor and smooth muscle myosin enhanced the originally ‘slow’ motor’s actin filament velocity, while the MLC2 that naturally assembles with the slow soleus or β-cardiac myosin slowed down the native fast myosin. It appeared that these specific MLC2 variants can increase or decrease the velocity depending on the myosin heavy chain isoform they are attached to. The changes in different kinetic and mechanical parameters of chimeric motors relative to the WT motor forms are listed in Table 1 and 2. In all but two chimeric motors i.e., S1s-MLC2B and S1f-MLC2v, the ‘t_on_’ that represents the duration of strong bound states correlated with the increase or decrease in the gliding velocity (Table 1 and 2). Although the trapping measurements were done at subsaturating ATP concentration, the relative changes in the ‘t_on_’ indicated changes in AM detachment rates among WT and chimeric motors. In comparison to its native forms, for chimeric motors, shorter and longer t_on_ relating to higher and lower rates of acto-myosin dissociation (1/t_on_), consequently resulted into faster or slower speed of actin filament speed, respectively. The powerstroke size was comparable between WT-S1f and S1f-chimeras and for WT-S1s and S1s-chimeras, except for S1s-MLCB. S1s-MLCB displayed similar t_on_ to S1s-CgmRLC that displayed ˃5 fold higher gliding speed. However, the other differences between the two chimeras included 3 fold shorter powerstroke size and slower rate of weak-strong transition (due to slow Pi release, Figure 3F and table 1) that may be responsible for the slower gliding speed for S1s-MLCB.

Typically, ADP release (strong bound state) rates are known to determine the gliding velocity for slow myosin. The solution kinetics measurement showed 2 phases of ADP release with marginal differences in the slow phase of ADP release rates for all slow myosin chimeras (Table 1). However, significant difference (upto ∼3 fold) in the ‘t_on_’ was observed in trapping kinetics measurements under subsaturating ATP concentration, and the gliding speeds varied by up to ∼10 fold among wild type and chimeric motors. A main difference between solution kinetics and single molecule trapping experiment is the load experienced by the motor. Thus, the observed differences in the parameters derived from the two methods could be explained by high sensitivity of the motors to the load as in trapping assays (even at the low applied values), and consequently, load sensitive ADP release influencing the mechanical output of the motors.

For the chimera, S1f-MLC2v, the reduction in actin filament gliding speed could be associated to the increase in binding affinity to actin, thereby, increasing the fraction of attached vs. detached motors, and generating a drag on gliding filaments as we previously reported [27]. We found that the dimeric chimera of fast heavy meromyosin, HMMf-MLC2v even displayed processive behaviour along actin filaments in single molecule studies.

We identified previously unrecognized aspect of the RLC’s task i.e., the effect on AM cross-bridge cycling by influencing the ATPase kinetics.

An obvious question raised by our findings is how diverse RLCs attribute such a response on myosin function. This is a very intriguing, yet quite complex question, which would require structural investigations and perhaps additional computational approaches to dissect possible long-range allosteric communication between the RLC and motor domain.

The different isoforms of RLC are similar in structure as predicted from their aa sequences, in particular, in the divalent cation-binding sites and proximal phosphorylatable serines in striated muscles [10]. They have relatively low sequence similarity, with the light chains MLC2B/ MLC2f and MLC2v are 86 % similar, while the cgmRLC exhibiting only 67 % sequence similarity with the fast MLC2B/ MLC2f isoforms.

RLC is categorized as an EF-hand superfamily member, harboring helix-loop-helix structure and a cation (Ca^2+^ or Mg^2+^) binding site at the N-terminus. Figure S2 depicts the conserved eight predicted helices, which forms four EF-hand domains. The binding of Ca^2+^ to the RLC is reported to induce a conformational change to more open state whereby the overall helical content of the molecule increases which subsequently positively affects the cross-bridge kinetics [58]. In our measurements we have no Ca^+2^ but Mg^2+^ in the reactions buffers. Additionally, the aa sequence for the helix 1 and 2 regions are 100 % identical in fast and slow RLC while the loop that serves the binding site for divalent cation has 3 aa difference. With the current knowledge however it’s challenging to foresee which specific feature/s might define particular RLCs modulating characteristic. Detailed structural and functional investigations would be required to address the diverse impact of the regulatory light chains.

This finding elucidates the specificity of RLCs towards distinct MHCs. In addition to understanding the fundamental role, precise details of modulatory role of RLC isoform is of particular clinical relevance as single point mutations in the human myosin heavy (β-MHC) and regulatory light chains (MLC2v) have been linked to causing Familial hypertrophic or dilated cardiomyopathy (HCM and DCM). Researchers have gained a plethora of valuable information not only from studies on human muscle tissue, but also from various animal models (e.g., mice and rabbit) and recombinant motor proteins about the motor dysfunction as a result of single point mutations. These studies often constituted motor components either from mixed isoforms (e.g., human mutant MLC2 with mice MHC and MLC1), or the mutations generated in different MHC isoform background (e.g., mouse α-MHC or Dictyostelium discoideum myosin II). Some studies were performed without light chain binding domains or motors with only MLC1 but not MLC2. Recent micro-mechanical characterization of human induced-pluripotent stem cell-derived cardiomyocytes myofibrils showed different (increased rates) kinetics of active force generation and relaxation to that of the adult human ventricular myofibrils. One possible explanation was that besides other protein isoform differences, the β-MHC expression did not fully match with the respective light chain expressions (i.e., MLC1v and MLC2v), rather atrial (MLC1a and MLC2a) light chains dominated over ventricle specific light chains [59].

Altogether, a striking revelation from our studies point that the complete ‘holoenzyme’ with the respective heavy and light chains is critical to understand the motor function and therefore should be taken into account in structure-function analyses.

Overall, our findings offer a highly compelling evidence of RLCs role in modulating myosin motor function by influencing the actomyosin ATPase kinetics. The functional differences among the myosin II family members appear to be only partly defined by the myosin heavy chain alone. Here, we show that variations in light chain composition crucially determine the mechanical output of a specific myosin II isoforms that is commonly found in different muscle types and various tissues. The importance of these finding extends beyond basic understanding of the myosin function. The study opens new perspectives to investigate the physiological relevance of light-chain mediated fine-tuning of myosin function in the context of muscle contraction and additionally for other classes of myosin involved in transport processes.

## Acknowledgments

### General

We thank and dedicate this work to late Prof. Bernhard Brenner. We thank Dr. Atsuko Iwane for a kind gift of BDTC-chicken gizzard RLC construct to Dr. Walter Steffen. We thank Petra Uta and Stefanie Nedel for the technical assistance in protein production and purification, Prof. Theresia Kraft and Bogdan Iorga for their critical comments on the manuscript, Dr Ante Radocaj for his comments on statistical analysis.

### Funding

This research was supported by a grant from Deutsche Forschungsgemeinschaft (DFG) to MA (AM/507/1-1). TW is supported by a grant from Fritz Thyssen Stiftung to MA (10.19.1.009MN). WS was supported in part by a DFG grant (STE 1697/1-2). GT was supported by the DFG grant TS 169/5-1.

### Author contributions

MA conceived the project and designed the experiments with inputs from AN. AN performed cloning and protein preparations. AN and MA performed motility experiments and data analysis. TW, MA and WS performed optical trapping measurements and analyzed the data. PF, IC, GT, and MA performed solution kinetics experiments. Solution kinetics data was interpreted by PF, GT and MA. Motility and single molecule data was interpreted by AN, TW and MA. MA wrote and edited the manuscript with assistance from AN, GT and the other authors. AN and TW contributed equally to this work.

