## Supplementary material for "Single molecule analysis reveals the role of regulatory light chains in fine-tuning skeletal myosin-II function": AM_Supplementary Information

**This file includes:**

(1) Figures S1 to S2:

Figure. S1. Gel images displaying RLC exchange

Figure. S2. Sequence alignment

(2) Captions for movies S1 to S2

Movie S1. Actin filament gliding for native fast (S1f) vs. slow myosin S1 (S1s)

Movie S2. Actin filament gliding for chimeric motors S1s- cgmRLC vs.S1f- MLC2v

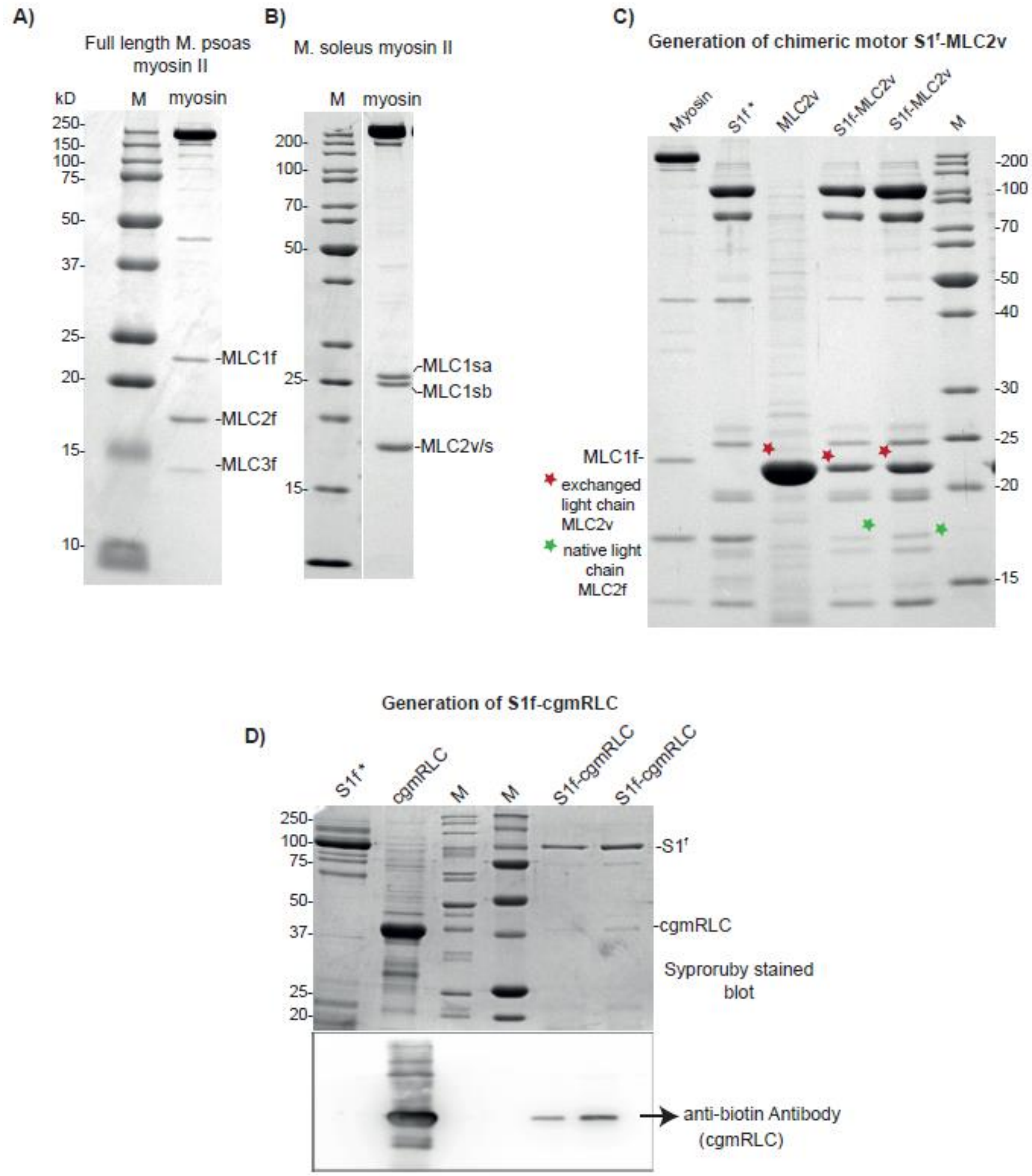


**Figure S1. *In vitro* reconstitution of myosin complexes.**

Gel images display the replacement of native RLC with recombinant RLC. Some examples are shown here. A) and B) Native full length myosin extracted from Rabbit M. Psoas and M. soleus muscle fibers. M. Psoas myosin II lane displays the 2- native ELC isoforms, a long one (~21 kDa, MLC1f) and a short one (~16 kDa, MLC3f). M. soleus myosin II lane also display 2-

distinct ELC isoforms (MLC1sa and MLC1 sb with molecular weight of 27 and 24 kDa, respectively), and one RLC isoform, MLC2v/s. Please note that in Figure B, the two lanes are taken from the same gel, but the middle lanes which are not relevant here were cropped out, as indicated by the space between the two lanes. C) S1f-MLC2v chimera generation, i.e., combination of fast myosin heavy chain with the slow RLC (MLC2v). Coomassie blue stained gel shows the replacement of native RLC with recombinant MLC2v. M- Marker in different gel images. Myosin - full length myosin, S1f is papain digested subfragment-1(S1). * indicates the S1 without actin purification step and therefore contains other digested products. Typically the S1 is purified by actin-cosedimenation assay followed by ATP induced release of functional myosin heads. D) Generation of chimeric form of motor with cgmRLC. Upper blot was syproruby stained, and the same blot was further probed with anti-biotin antibody to ensure the correct size and specificity of the relevant cgmRLC band. CgmRLC was biotinylated. S1f-cgmRLC - 2 different concentrations run on the last 2 lanes. Please note that only coomassie stained gels were used for densitometric analysis to measure the exchange efficiency for different preparations of protein exchange. Syproruby stained blots cannot be used for densitometric analysis as the band transfer efficiency from the gel to the membrane for proteins with different molecular weights cannot be efficiently controlled i.e., the high molecular weight bands require longer transfer time from gel to the nitrocellulose membrane while the low molecular weight proteins needs shorter transfer time. Therefore, the optimal transfer could not be attained for different size. The identity of the cgmRLC bands was checked with the western blot using anti-biotin antibody (note that biotinylated cgmRLC samples used for the gels) while Coomassie stained gels were used for quantification. Marker molecular weight bands are indicated as M in the left image panel.

```
Q7M2V4     MSPKKA-----KKRA--EGANSNVFSMFEQTQIQEFKEAFTIMDQNRDGFIDKNDLRDTF
P10916     MAPKKA-----KKRA--GGANSNVFSMFEQTQIQEFKEAFTIMDQNRDGFIDKNDLRDTF
P02608     MAPKKA-----KRRAAAEGGSSNVFSMFDQTQIQEFKEAFTVIDQNRDGIIDKEDLRDTF
Q96A32     MAPKRA-----KRRTV-EGGSSSVFSMFDQTQIQEFKEAFTVIDQNRDGIIDKEDLRDTF
P02612     MSSKRAKAKTTKKRP--QRATSNVFAMFDQSQIQEFKEAFNMIDQNRDGFIDKEDLHDML

Q7M2V4     AALGRVNVKNEEIDEMIKEAPGPINFTVFLTMFGEKLKGADPEETILNAFKVFDPEGKGV
P10916     AALGRVNVKNEEIDEMIKEAPGPINFTVFLTMFGEKLKGADPEETILNAFKVFDPEGKGV
P02608     AAMGRLNVKNEELDAMMKEASGPINFTVFLTMFGEKLKGADPEDVITGAFKVLDPEGKGT
Q96A32     AAMGRLNVKNEELDAMMKEASGPINFTVFLTMFGEKLKGADPEDVITGAFKVLDPEGKGT
P02612     ASMGK-NPTDEYLEGMMSEAPGPINFTMFLTMFGEKLNGTDPEDVIRNAFACFDEEASGF

Q7M2V4     LKADYVREMLTTQAERFSKDEIDQMFAAFPPDVTGNLDYKNLVHIITHGEEK---
P10916     LKADYVREMLTTQAERFSKEEVDQMFAAFPPDVTGNLDYKNLVHIITHGEEKD--
P02608     IKKQFLEELLTTQCDRFSQEEIKNMWAAFPPDVGGNVDYKNICYVITHGDAKDQE
Q96A32     IKKKFLEELLTTQCDRFSQEEIKNMWAAFPPDVGGNVDYKNICYVITHGDAKDQE
P02612     IHEDHLRELLTTMGDRFTDEEVDEMYREAPIDKKGNFNYVEFTRILKHGAKDKDD
```

**Figure S2. Sequence alignment**

Regulatory light chain sequences were aligned with CLUSTAL W multiple align tool by Kalign (2.0); Q7M2V4: Rabbit MLC2v, P10916: Human MLC2v, P02608: Rabbit MLC2B, Q96A32: Human MLC2B, P02612: Chicken cgmRLC. The sequences marked with grey background represent the 8 helical structures. Fast, MLC2B from human and rabbit shares 98% sequence similarity, so does the slow MLC2v from both species. The non-identical amino acids in fast and slow RLC isoforms are indicated in red. Blue arrows point at the 3 amino acids different in the divalent cation binding loop of fast (MLC2B) and slow (MLC2v) RLCs. The cgmRLC shares low sequence homology with both slow and fast MLC2, the non-identical amino acids indicated in bold black letters.

**Movie legends.**

**Movie S1:** *Actin filament gliding for native fast (S1f ) vs. slow myosin S1 (S1s)*

The *in vitro* actin filament gliding on myosin WT-S1f (left panel) and WT-S1s (right panel) coated surface at saturating ATP concentration and 22 °C. Data were acquired at 5 frames /s, scale bar 10 µm; Movie was sped up to run at 20 frames/s.

**Movie S2:** *Actin filament gliding for chimeric motors S1s- cgmRLC vs.S1f- MLC2v*

The *in vitro* actin filament gliding on S1s- cgmRLC (left panel) and S1f- MLC2v (right panel). Fast and slow motors were indistinguishable after the RLC swapping. Data were acquired at 5 frames /s, scale bar 10 µm; Movie was run at 20 frames/s.
